# Combining Low toxic dose Tramadol and smoking is relatively safe unless you stop them: An Animal model evidence of Endoplasmic reticulum stress

**DOI:** 10.1101/2023.05.25.542154

**Authors:** Doaa Ghorab, Ejlal M. Abuelrub, Mohamed Hamdi Gharaibeh, Alaa Yehya, Ramada R Khasawneh, Laila M Matalgah, Ahmed Mohamed Helaly

## Abstract

Low toxic doses of tramadol induced animal brain cortex apoptosis and hippocampus injury. Adding nicotine reverted hippocampus pathological changes without triggering brain injury. The expression of CHOP protein in real-time PCR showed mild Endoplasmic reticulum stress (ER) in rats’ brains. Histological, immunohistochemical, and western blotting analysis of CHOP and BIP chaperones demonstrated Endoplasmic reticulum stress in brain and liver tissue samples. Furthermore, the levels of apoptosis and autophagy markers demonstrated a mild increase. Adding Nicotine relatively decreasedbrain and liver ER stress. The combined profile was considerably protective in comparison to administering each drug separately. Mild ER stress is essential for normal cell functions. The blood level of serotonin was high in all study groups with a marked increase in its level when tramadol and nicotine were combined. Low toxic doses of tramadol in combination with nicotine were safe at the reproductive system level which was evaluated by histological examination and animal blood androgen assay. Generally, combining low-dose tramadol with smoking was found to be safe in various animal tissues and organs, however, the high serotonin level in the blood can be critical and associated with a high risk of serious withdrawal and pathological consequences. Serotonin receptor blockers such as olanzapine may increase systemic serotonin levels and need further investigation to utterly pinpoint their roles in managing mood disorders.

## Introduction

Tramadol is a centrally-acting analgesic drug that is prescribed to treat moderate to severe acute or chronic pain **(Minami, Ogata, and Uezono 2015)**. Nicotine, a lipid-soluble alkaloid, is one of the most accessible and abused chemicals worldwide **(Wonnacott, Sidhpura, and Balfour 2005; Marcon et al. 2018)**. Co-administration of cigarettes and opiates has a cumulative impact on inducing toxicity and health-related complications of many body systems **(Hursh et al. 2005)** resulting in excessively high rates of morbidity and mortality among regular drug users **(Khademi et al. 2012; Nalini et al. 2021)**. Smokers are more likely to experience chronic pain of greater intensity, a greater number of painful sites, more associated impairments **(Yoon et al. 2015)**, and negative impacts on occupational and social functioning **(Hooten, Townsend, Bruce, Shi, et al. 2009)**. Compared to non-smokers, they have higher pain scores and greater demand for opioids during and after surgery **(Sandén, Försth, and Michaëlsson 2011).** Opiates use especially prescribed replacement medicines, has been reported to encourage smoking habits **(Chun et al. 2009)**. A previous study which included a sample of 48 smokers who are also tramadol addict patients, suggested that Tramadol increased the severity of nicotine dependence, as the mean Fagerstrom Test for Nicotine Dependence (FTND) score dropped from 6.67 during the tramadol addiction phase to 4.31 five weeks after stopping using tramadol **(Shalaby, Sweilum, and Ads 2015)**. in addition,, studies have indicated that smokers are more likely than nonsmokers to be on opioid pain medications dependent for longer periods and at greater dosages **(Novy et al. 2012; Hooten, Townsend, Bruce, Warner, et al. 2009)**. Higher levels of reported pain among smokers may also be associated with higher levels of depression, anxiety, and sadness **(Zale, Maisto, and Ditre 2016)**.

In an attempt to explore the molecular underpinnings of nicotine and opioid addiction, several experimental studies were conducted. **Azmy et al, 2018** reported that nicotine augmented oxidative stress, inflammatory reaction, and possibly apoptosis post-combined exposure to tramadol/nicotine in brain samples of mice. Tramadol abusers were reported to suffer from disrupted fertility profiles **(Azmy, Abd El-Rahman, et al. 2018).** In parallel, previous studies reported that nicotine has a dose-dependent detrimental impact on sperm characteristics **(Kim et al. 2008).** Recently, it was found to induce endoplasmic reticulum stress in the placenta in the rat model **(Wong, Holloway, and Hardy 2016b).** On the other hand, recent experimental work demonstrated the potential protective mechanism of nicotine on injured nerve cells **(Zhao et al. 2020).** Despite the induction of oxidative stress, nicotine exerts anti-inflammatory action mediated by alpha 7 nicotine receptors **(Mabley, Gordon, and Pacher 2011; Hajiasgharzadeh et al. 2020).** It was reported that nicotine decreased tumor necrosis alpha and interleukin 1 beta **(Liu et al. 2018).**

The endoplasmic reticulum is an essential cell component dealing with protein production, processing modifications, and transport. Endoplasmic reticulum stress is regarded as a physiological process in addition to pathological conditions. If the cell is subjected to an endoplasmic reticulum load, it stimulates the chaperone battery to protect the cell against injury. If the condition is overwhelming, the cell stimulates an apoptosis reaction. BIP(GRP78), an hsp70-related, endoplasmic reticulum-resident protein, and CHOP (CCAAT-enhancer-binding protein homologous protein) proteins are chaperones expressed to protect the endoplasmic reticulum from strain **(Bengesser et al. 2018).**

As a result, nicotine experimentally might have contradictory effects on the brain. In light of these outcomes, this work aims to investigate the possible modulatory effects of nicotine on tramadol-induced neuropathological changes in the brain and testicle tissues. In addition, it evaluates the expression of specific markers of endoplasmic reticulum stress post-combined exposure to nicotine and Tramadol.

## Material and Methods

Animal Groups: 6 animal groups were selected for the current study. The animals were 6-week-old Sprague Dowley rats with 150 to 200 mg weight. The course of the study was a 3-week experiment and the animals were injected 5 times per week. The experiment was done in the animal house of the Jordanian University of Science and Technology Lab. The handling of the animals has been done according to the ethical committee review that accepted the project by code number (752/ 12/4/86).

1- Group 1 Tramadol 20mg/ kg per animal by oral gavage.
2- group 2 10 mg/kg tramadol per animal by oral gavage.
3- group 3 combined tramadol/nicotine 20 mg tramadol/ 125 mcg/kg nicotine per animal.
4- group 4 nicotine 125mcg/kg s.c.
5- olanzapine 3mg/kg per animal by oral gavage.
6- Negative control oral saline.

## Methods

### Hematoxylin and Eosin

**Firstly,** paraffin sections weredewaxed in xylol and hydratedthrough descending grades of alcohol to distilled water. Then, sections were put in Harris hematoxylin for 5 minutesand washedin running tap water for 5 minutes. The next stage was a differentiation of the tissue sections in 1% acid alcohol (1% HCl in 70% alcohol) for 5-10 seconds. Sections were washed in running tap water again until they became blue. Then, staining in 1% eosin for 10 minutes was carriedout and was followed by washing with distilled water. Finally, the sections were dehydrated through ascending grades of alcohol, were cleared in xylol, and were mounted in Canada balsam **(Lyle 1947).**

### Immunohistochemistry

Slides underwent deparaffinization by incubation in the oven at 62-65 C for 30 mins, then they were incubated with xylene anddehydrated indescending grades of alcohol(100%-90%-70%) each for 5 min. The slides were washed in a buffer solution for 10 min. The next step was epitope retrieval by boiling in a pressure cooker with 0.01 M HIER Citrate Buffer pH 6.5. Blocking of endogenous peroxidase using the 3% hydrogen peroxide in methanol for 5 min was applied. The samples were then washed in the buffer for 5 min, thenthe slides were incubatedwith proteinase K 0.04% for 5 min. After that, the slides were washed with PBS for 5 minutes and incubated with rabbit CHOP polyclonal antibody (MyBioSource catalog no, MBS126028) with a dilution of 1:500 at room temperature in a humid chamber for 2 hours, and a Mouse Caspase-8 monoclonal antibody (MyBioSource catalog no, MBS8808615) with a dilution of 1:200 overnight and rabbit polyclonal P53BP1(GenoChem catalog no. GW2740R) with a dilution of 1:200 overnight were used. The slides werewashed in PBS 3 times for 2 min each, and then 2 drops of the secondary antibody were added and incubated for 10 min. The next step was washing PBS 3 times for 2 min each. Two drops of streptavidin-biotinwereadded for 30 min at room temperature andwashedin PBS 3 times for 2 min each. Diaminobenzidine (DAB) with a dilution of 1:50 was added and used as the chromogen for 1-3 min (Abcam rabbit specific HRP/DAB detection kit ab64261). Again, the slides were rewashed in PBS 3 times for 2 min each, and 2 drops of Mayer hematoxylin counterstain will be added for 1-3 min. The slides were put in the buffer for 30 seconds, then were washed in distilled water, dehydrated using ascending grades of alcohol, and cleared in xylene for 5 min before mounting **(Hicks et al. 2017).**

### Western blot

The protein levels of BIP (Cat# MBS857422, Mybiosource, USA), CHOP (Cat# MBS126028, Mybiosource, USA), Caspase 8 (Cat# MBS8808615, Mybiosource, USA), and LCIII (Cat# MBS2520564, Mybiosource, USA) were measured by western blotting using species-specific antibodies. Briefly, total protein levels were measured by NanoDrop™ Lite Spectrophotometer (ThermoFisher Scientific) and 50 μg of protein was loaded onto SDS-PAGE. After electrophoresis, proteins were transferred to the PVDF membrane and incubated with primary antibodies and corresponding secondary antibodies. The membranes were visualized using a Gel documentation imaging system (Vilber, France) and bands were quantified using ImageJ software for densitometry.

## Real-Time PCR

### -RNA Isolation

Total RNA was extracted using (Jena Bioscience, Germany) extraction kit. The steps of eluting the RNA were applied to the manual.

### cDNA Synthesis

cDNA synthesis kit, which was used in the study, was EasyScript First Strand cDNA Synthesis Supermix Cat. No. AE301

### Real-time Polymerase Chain Reaction (PCR)

To determine the expression of the target genes, relative quantitative real-time PCR will be performed using SYBR-green HOT FIREPOOL EVAGREEN q PCR Supermix from SOLIS BIODYNE. The qPCR cycle protocol is used according to the following:

**Table 1.** illustrates the cycle of real-time PCR for CHOP, BIP, and GAPDH gene expressions

|  |  |  |
| --- | --- | --- |
| Initial activation | 95 degrees Celsius | 12 min |
| Denaturation | 95 degrees Celsius | 15 sec |
| Annealing | 62 degrees | 30 sec |
| Extension | 72 degrees | 30 sec |

The initial activation is done once, and the remaining component of the cycle is repeated 40 times. The results were evaluated by the BIO-RAD CFX maestro program. The samples were reevaluated in duplicate to confirm the results.

**Table 2.**
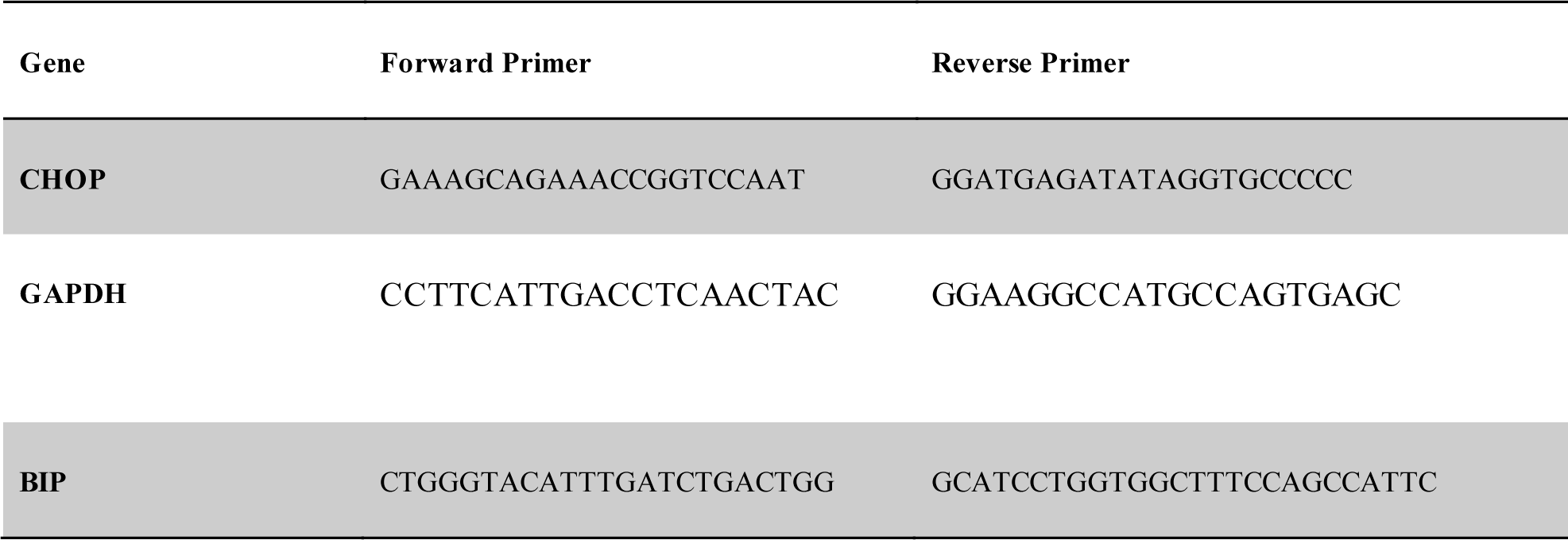
describes the primers of CHOP, GABDH, and BIP for qPCR reaction:

### 5-ELIZA technique

The serum samples have been frozen at minus 80 degrees till examination for ELIZA. The samples were tested according to the kit’s manual. The methods were obtained from the kits CatLog numbers: ELK8954 for Rat serotonin, ELK6024 for Rat androgen, and ELK8953 for Rat Dopamine kit. All kits were obtained from ELK Biotechnology Company.

### Statistical analysis

Statistical analyses of data were performed using the GraphPad Prism version 9.0 for Windows, (San Diego, California). The P values were determined using a one-way ANOVA Test. The differences were considered significant if P values were 0.05. A test has been applied for Figure number 8.

## Results

Histological and immunohistochemical results: Histological examination has been conducted on different animal groups: G1 high tramadol dose (20mg/kg day), G2 low tramadol dose (10mg/kg/day), G3 mixed nicotine /high tramadol, G4 nicotine group 125mcg/kg/day, G5 olanzapine 3mg/kg dose (low toxic dose, and G6 saline negative control group.

Mild inflammation was observed in the high-dose tramadol group and the Combined tramadol/nicotine-treated group (Figure 1). The combination neither did improve the mild toxic profile nor aggravated it. Lower tramadol, olanzapine, and the negative control were safe.

**figure (1):**
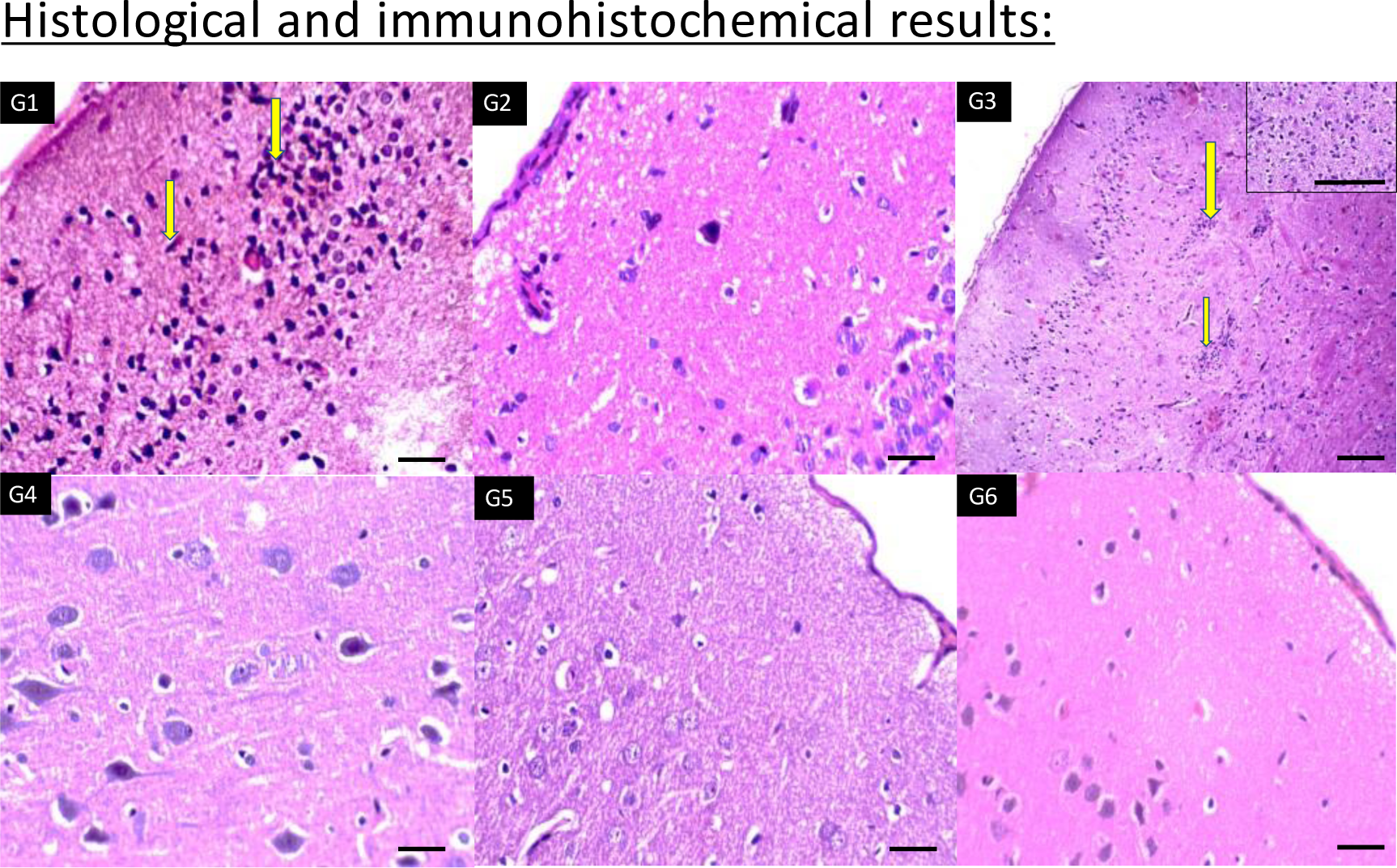
Sections of brain cortex of six groups showing increased aggregates of inflammatory cells in cortex of G1 (high dose tramadol) and G3 (Nicotine+tramadol) (Yellow arrows) (x400,400,100,400,400,400 respectively)

In Figure 2 the histological findings showed injured neurons in the low tramadol group with vasodilatation in the higher tramadol group. The Nicotine profile showed healthy hippocampus.

**figure (2).**
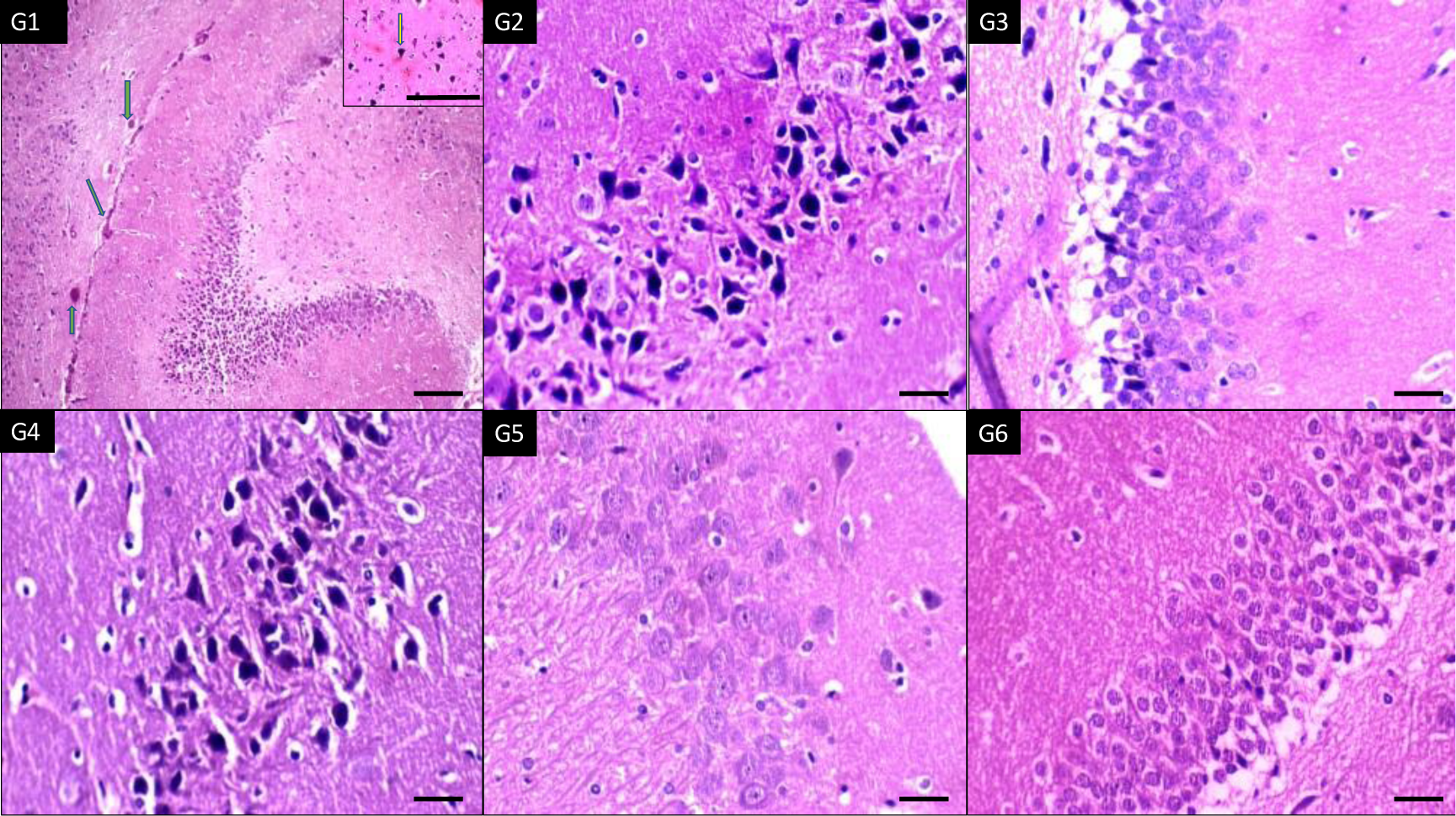
sections of brain hippocampus of six groups showing dilated congested blood vessels in G1(high dose tramadol) (green arrows) and increased degenerated neural cells in G2 (low dose tramadol) and (nicotine+tramadol) (X100, 400,400,400,400,400 respectively).

In Figure 3; the histology showed injured white matter in all nicotine, and tramadol groups. Again, high vascularity was demonstrated in higher tramadol dosage.

**figure (3):**
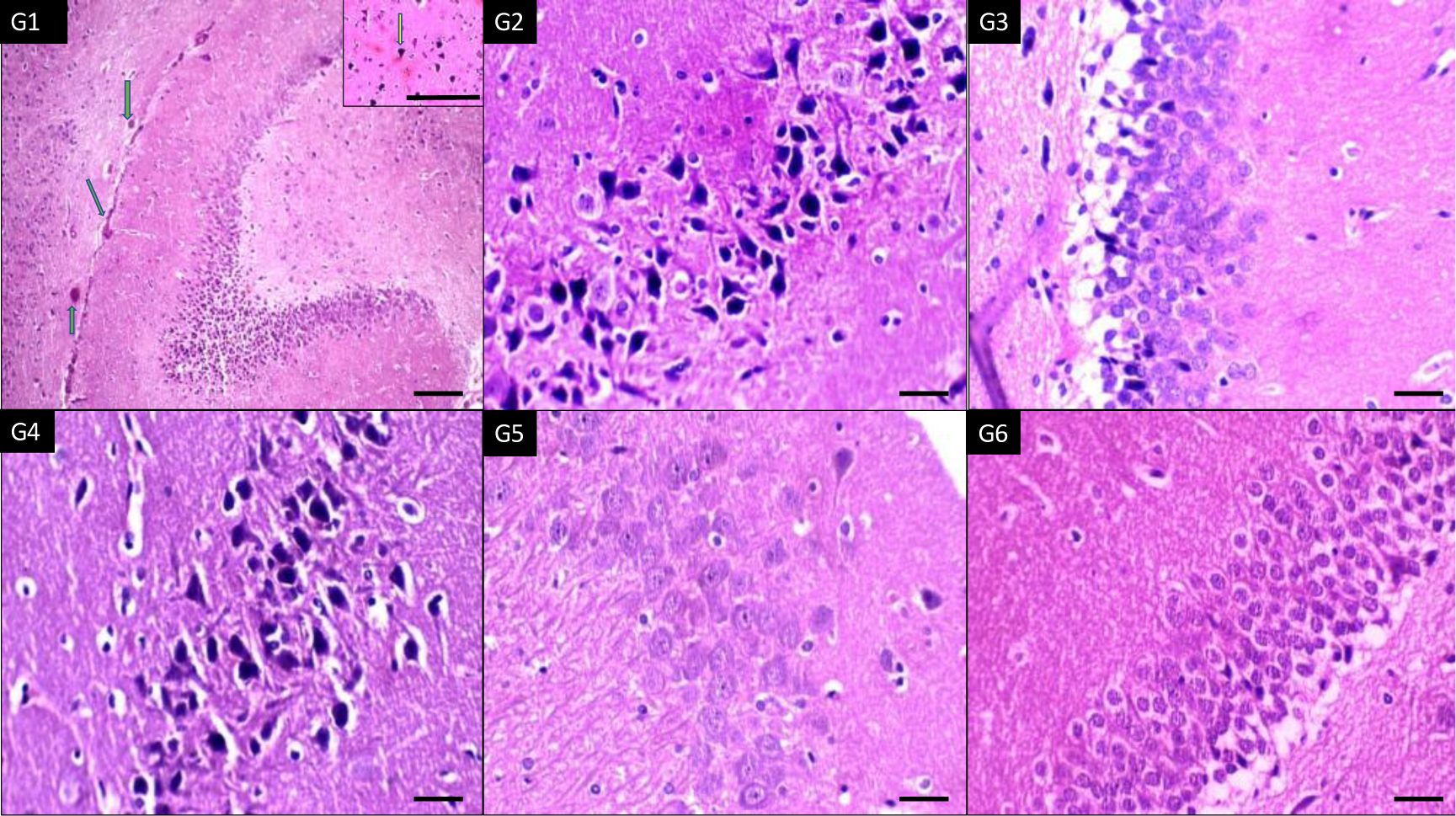
sections of brain white matter of six groups showing dilated congested vessel in G1(high dose tramadol) (green arrow) and increased inflammatory microglia in all groups in comparison to G6 (normal control) (yellow arrows) (x400 all)

In Figure 4 the samples were stained with CHOP immunohistochemical stains where the six groups demonstrated positive staining with marked reaction in the combined nicotine/tramadol group. Another immune staining with p53 showed higher expression in the combined nicotine/ tramadol group. On staining the animals’ brain cortexes with p53, the opposite was detected where the lower expression has been demonstrated in the nicotine-related groups.

**Figure (4):**
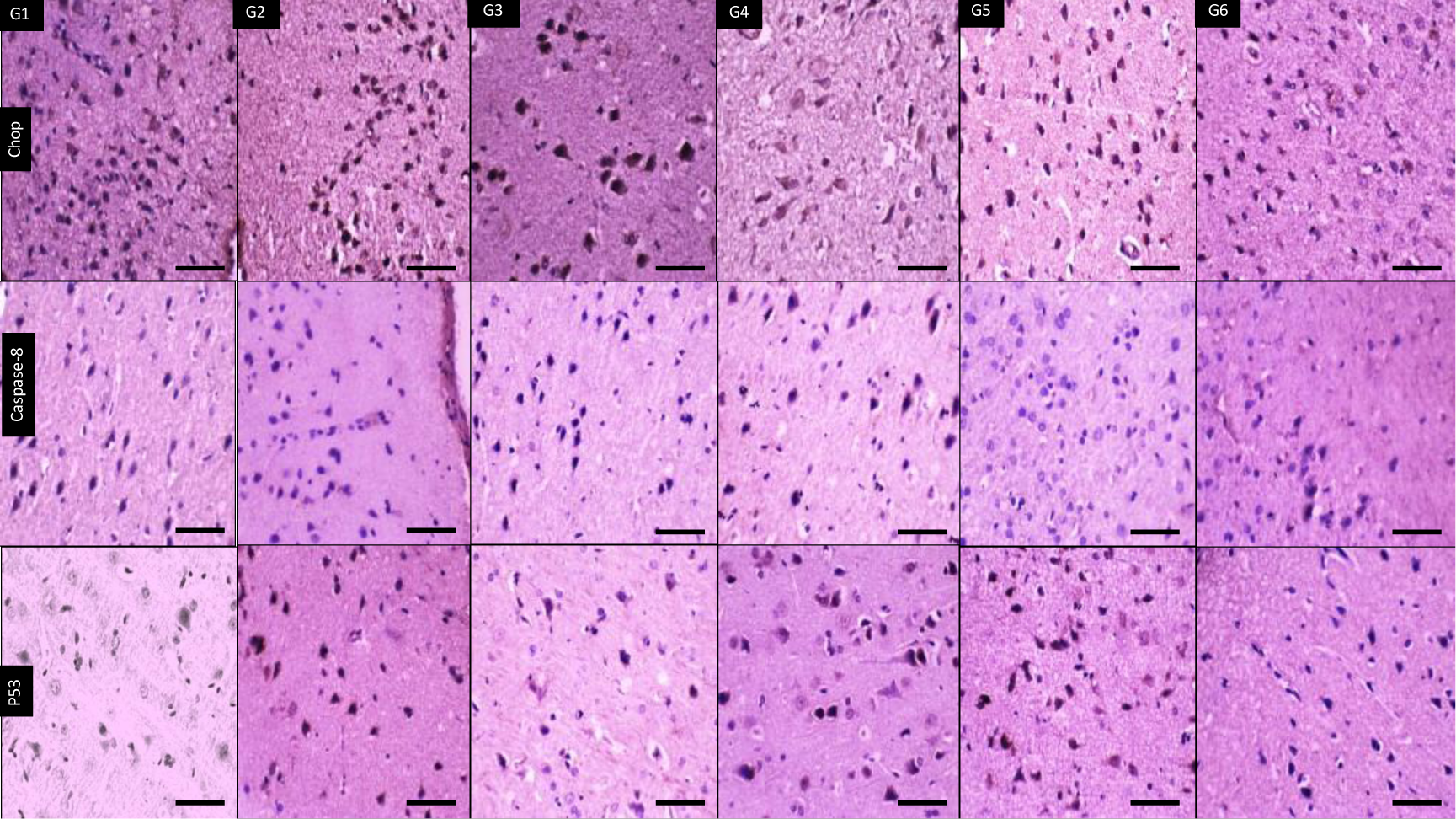
Immunohistochemical staining sections of brain cortex of six groups showing positive cytoplasmic expression of Chop antibodies in all groups, including G6(normal control) with strongest expression in G3 (nicotine+ tramadol). Negative expression of caspase-8 antibodies in all groups except G4 (nicotine) with mild cytoplasmic positivity. P53 is expressed in all groups with minimal positivity in G3 (nicotine+tramadol) and G6 (normal control). (X400 all)

In Figure 5, staining the animal hippocampus showed the highest reaction in the nicotine group than the combined nicotine/ tramadol. Lower immune staining was recorded in the other groups. Caspase 8 expression was active in all groups with less focal reaction in nicotine/tramadol animals. P53 immune stain showed minimal activity in nicotine groups.

**figure (5):**
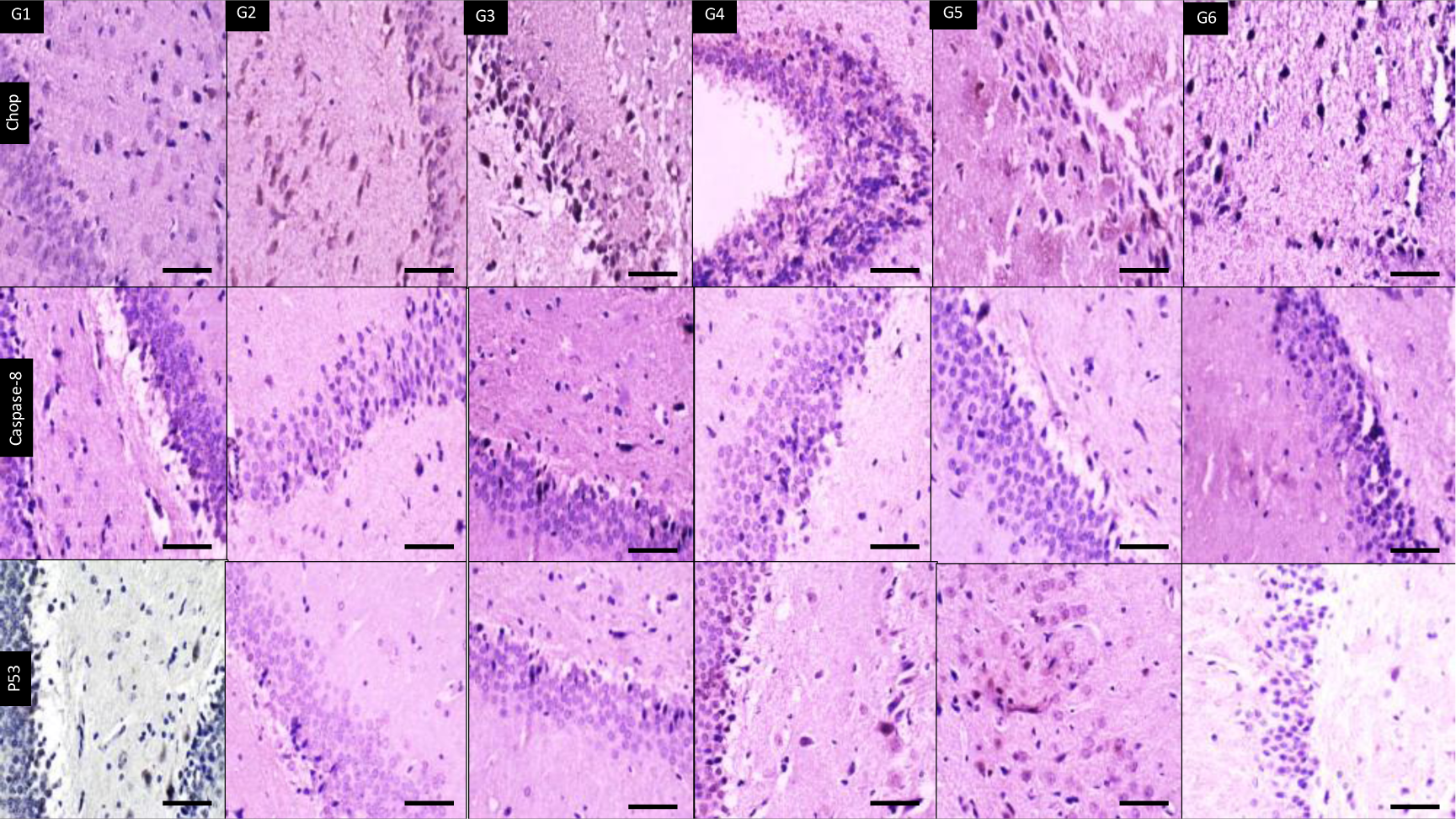
Immunohistochemical staining sections of brain hippocampus of six groups showing different expression of CHOP antibodies with maximum expression in G4 (nicotine) followed by G3 (nicotine+tramadol), then G2 (low dose tramadol) and G5 (olanzapine) with minimal expression in G1 (high dose tramadole), and G6 (normal control). Caspase-8 antibody is negative in all groups except in G3 (nicotine+tramadol) with focal positivity followed by G1 (high dose tramadol), focal positivity of caspase -8 in G2 (low dose tramadol) and G6 (normal control) is seen. P53 antibody is negative in all groups except showing minimal positivity in G4 (nicotine) and G3 (nicotine+tramadol). (X400 all)

In Figure 6 nicotine, tramadol, and olanzapine showed increased CHOP expression in the white matter with the highest stain in combined tramadol/nicotine.

**figure (6):**
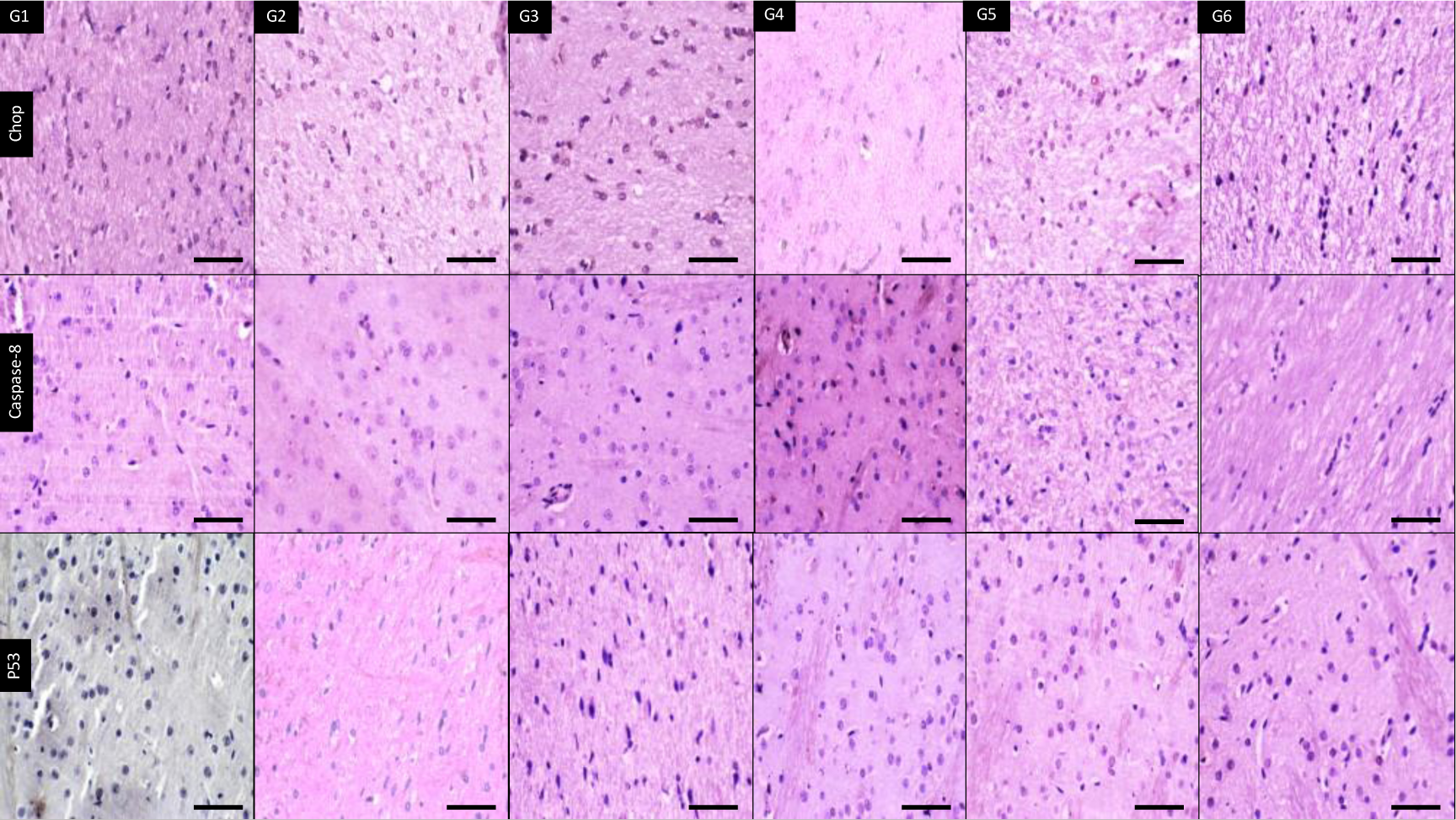
Immunohistochemical staining sections of brain white matter of six groups showing different expression of CHOP antibody with highest positivity in G3 (nicotine+tramadol) followed by G2 (low dose tramadol) and by G1 (high dose tramadol). Caspase-8 antibody is negative in all groups except minimally expressed in G4 (nicotine). P53 antibody is negative in all groups. (X400 all)

In Figure 7, the histological picture of liver cells was safe except for the vascular dilatation of the portal vein and mild hepatitis. The increase in the tramadol dose was associated with more vascularity. Nicotine addition was related to an increase in hepatitis, but the reaction was limited and focal.

**Figure (7):**
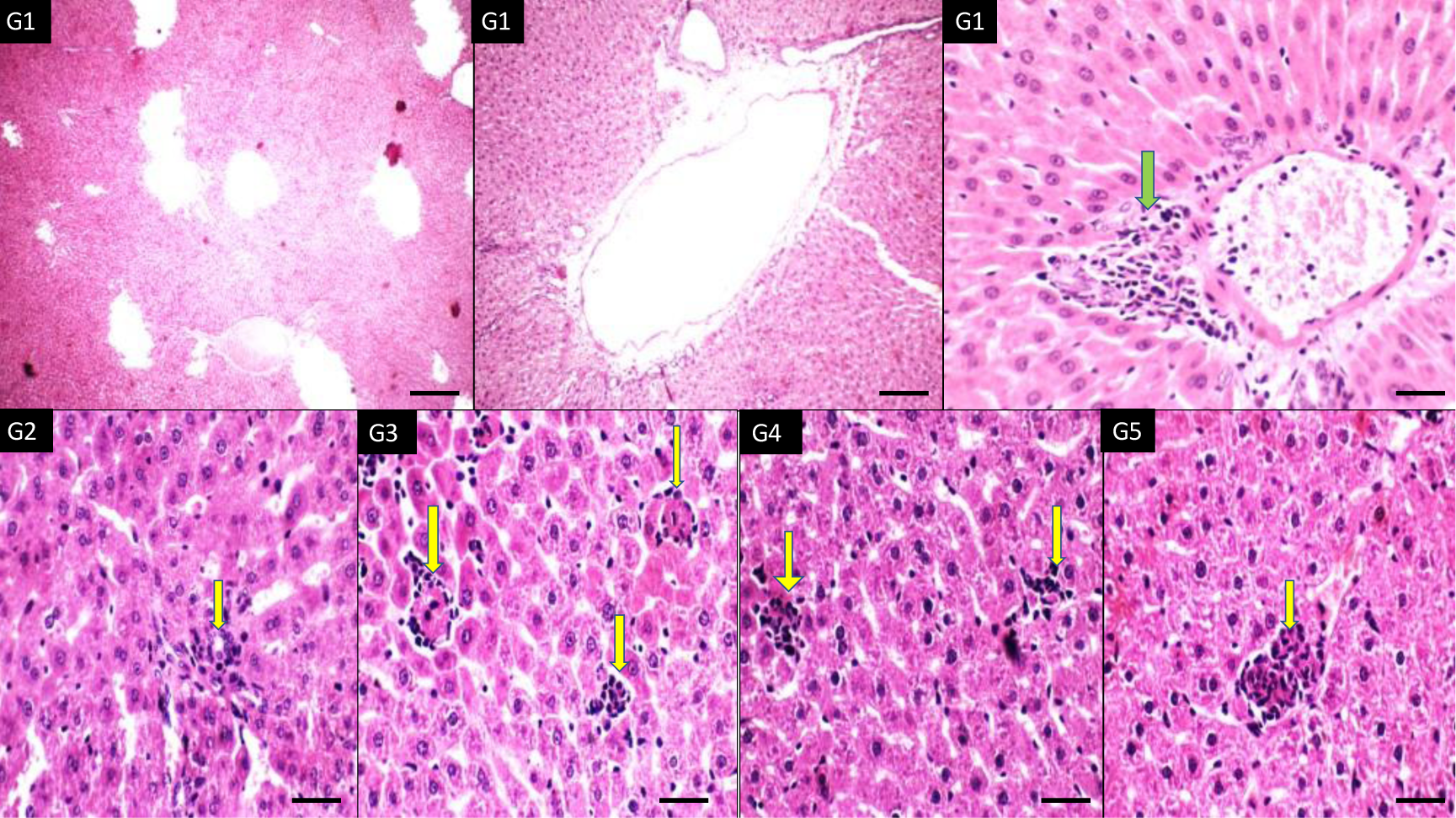
H&E sections from liver of six groups showing diffuse dilatation of portal veins and mild portal hepatitis (green arrow) in G1 (high dose tramadol) while all other groups show focal lobular hepatitis with more severity in G3 (nicotine+tramadol) and G4 (nicotine). (X400 all)

In Figure 8, the H&E staining showed normal spermatogenesis machinery in all groups except mild effects in high tramadol group. Focal Leydig cell proliferation was demonstrated with combined tramadol/nicotine.

**Figure (8):**
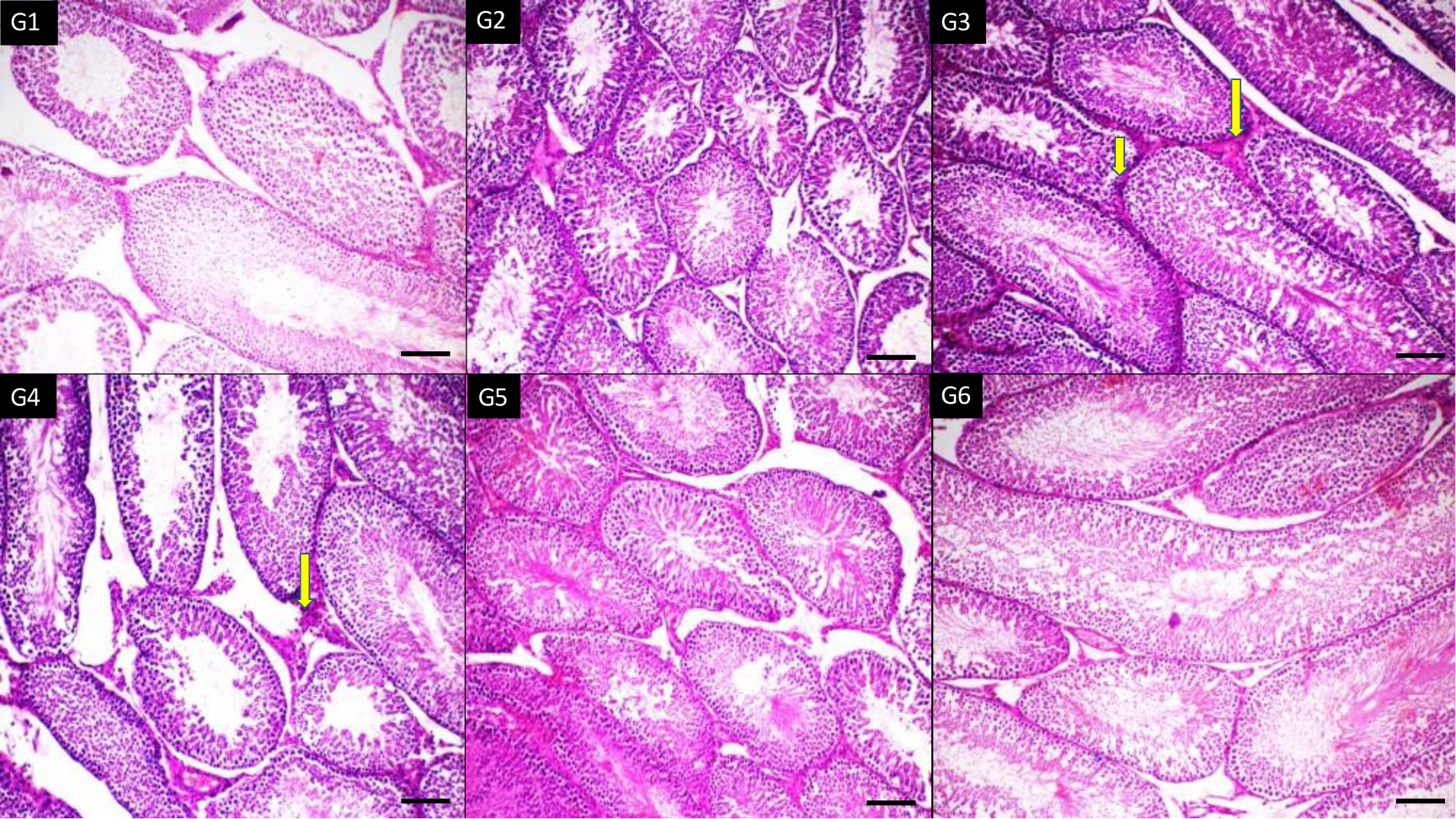
H&E sections of testis of six groups showing normal spermatogenesis in all groups (score 10) except G1 (high dose tramadol) show mild decreased spermatogenesis (score 9). There is focal Leydig cell hyperplasia in G3 (nicotine +tramadol) and in G4 (nicotine).(X400 all)

Figure 9 showed the western blot expression of CHOP chaperone as a marker of endoplasmic reticulum stress. The experiment tested 5 groups against the negative control group. The groups are 1-high tramadol (20 mg/kg group), 2-Low tramadol (10 mg/kg), combined high tramadol/nicotine group, 4-nicotine group 125 mcg/ group, 5-olanzapine 3mg/kg group, 6-saline negative control group. The highest protein expression was in the nicotine group. The negative control showed also a positive expression similar to a high tramadol dose. The low tramadol group expressed higher CHOP expression; combined nicotine/ tramadol expressed a better profile than nicotine alone.

**Figure (9):**
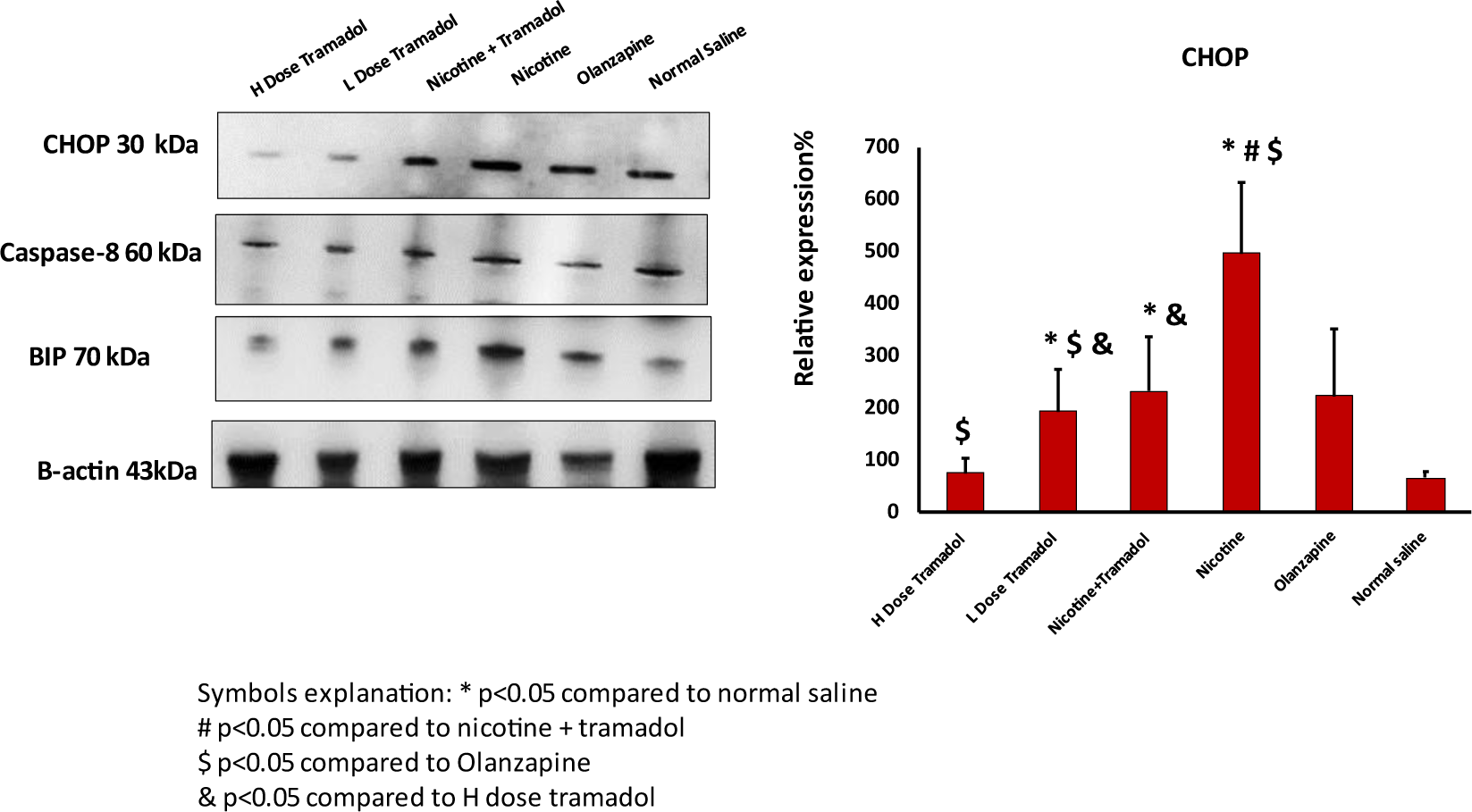
showed western blot protein expression of CHOP endoplasmic reticulum stress (ER stress) markers in rat brains. The figure also demonstrated the Caspase expression and the other BIP ER stress marker. The graph on the right side demonstrated the relative expression of Beta-actin protein. As regards CHOP expression, the highest expression was recorded in nicotine, then low tramadol and combined tramadol /nicotine that expressed less expression than the single nicotine category.

Figure 10 The BIP chaperone western blot results were supporting findings to the CHOP expression. Indeed, the negative control group has had a basal expression of BIP as well as CHOP. The most abundant expression was the nicotine group. The nicotine and combined groups expressed the two chaperones CHOP and BIP similarly. As regards caspase 8, the highest group was again the nicotine group and the lowest group was the negative control group. The other groups showed moderate protein expression.

**Figure (10):**
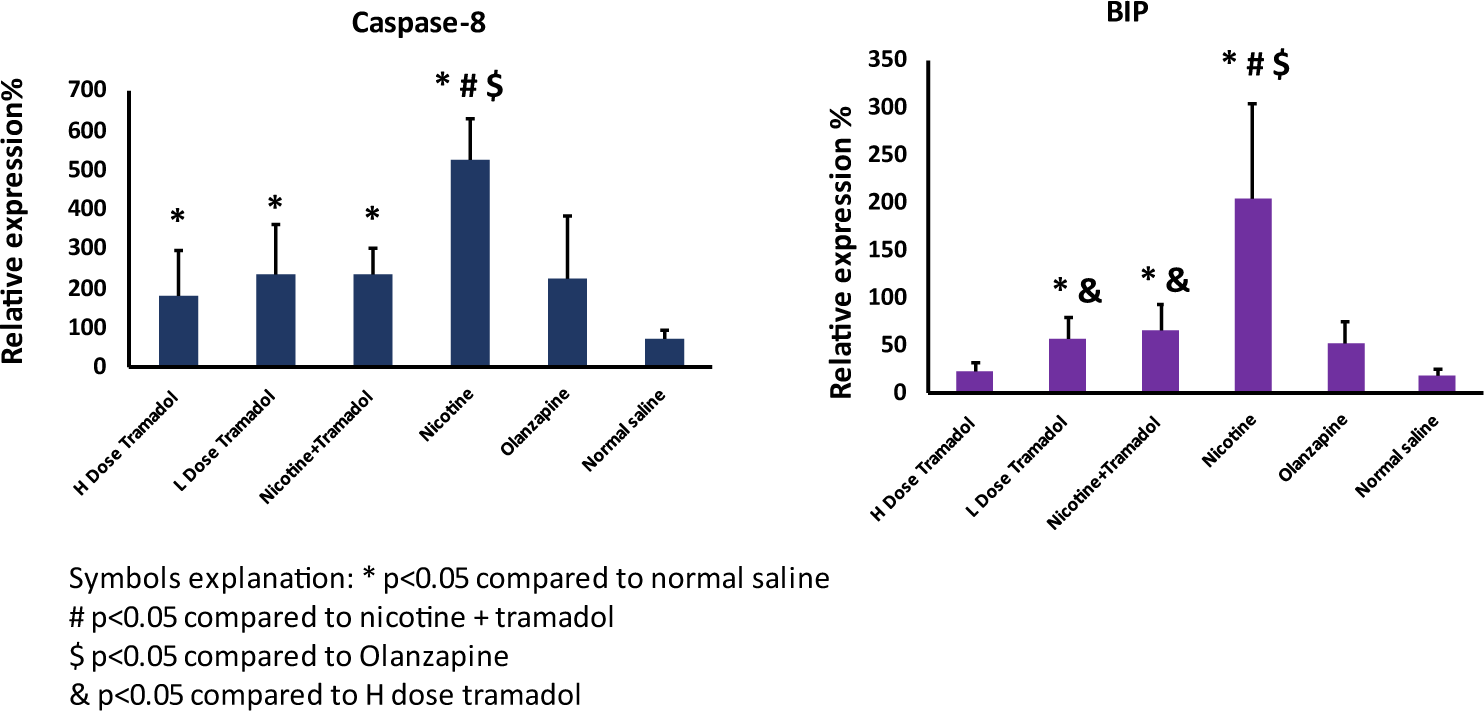
showed the quantitively expressed Caspase 8 and Bip chaperone. The highest expression of both was demonstrated in the nicotine group reflecting apoptosis and ER stress in rat brain samples.

Figure 11,12 showed western blot expressions of CHOP, BIP, and LCIII in the liver samples taken from the rats for the same groups as displayed in figure 1,2. The picture became more classic where the highest expression was as before in the nicotine group. The lowest expression was in the negative control group. The lower tramadol response was more favorable than the higher doses. The relative expressions of CHOP, BIP, and LCIII were in the same way. It can be noticed that the caspase 8 response expressed nearly the same pattern as the autophagy response detected by LCIII.

**Figure (11):**
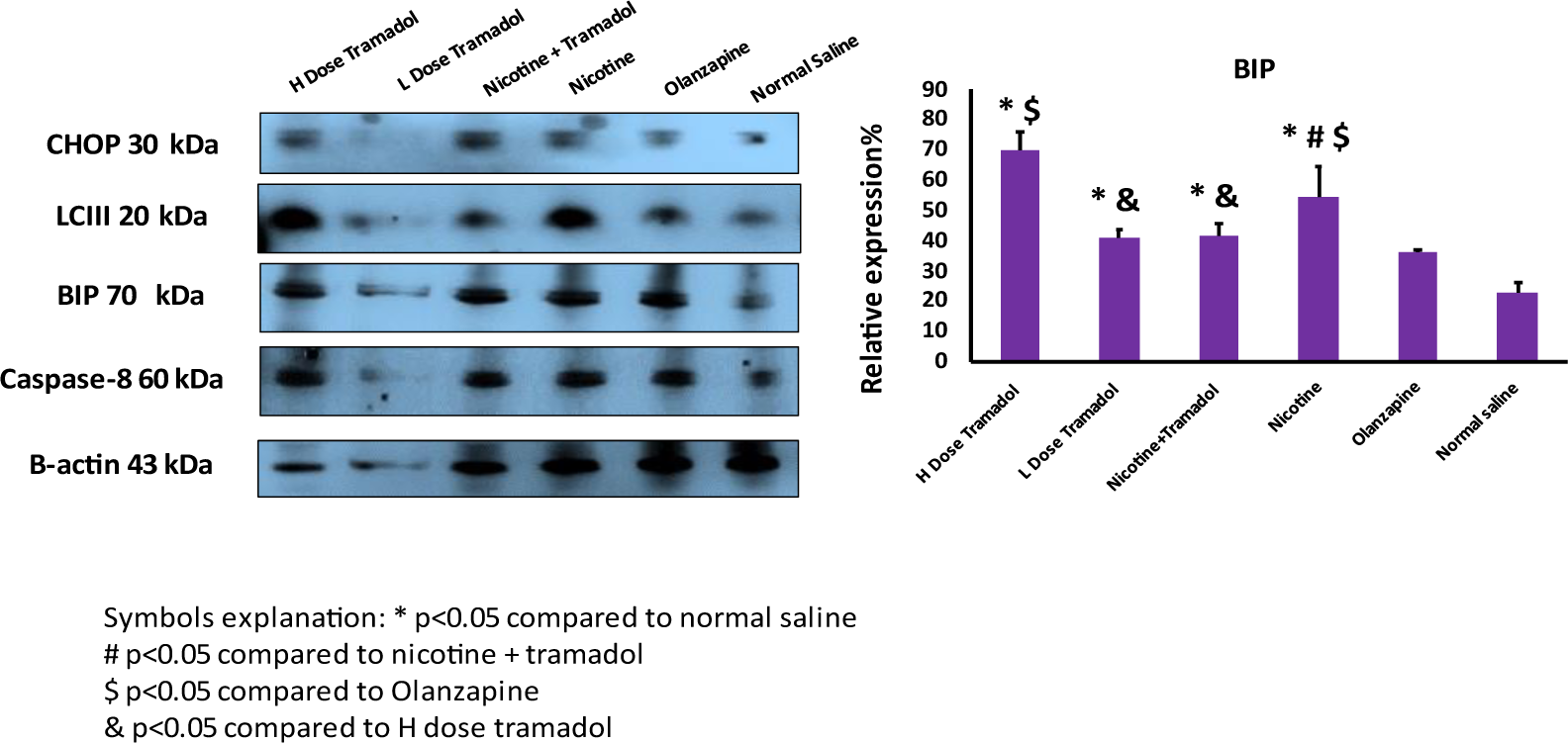
showed the western blot expression of CHOP, LCIII, BIP, and the control actin in the rat liver samples. The highest relative expression of ER stress marker was noticed in the tramadol and nicotine groups respectively. the best response was detected in the combined nicotine/ tramadol and low tramadol.

**Figure (12):**
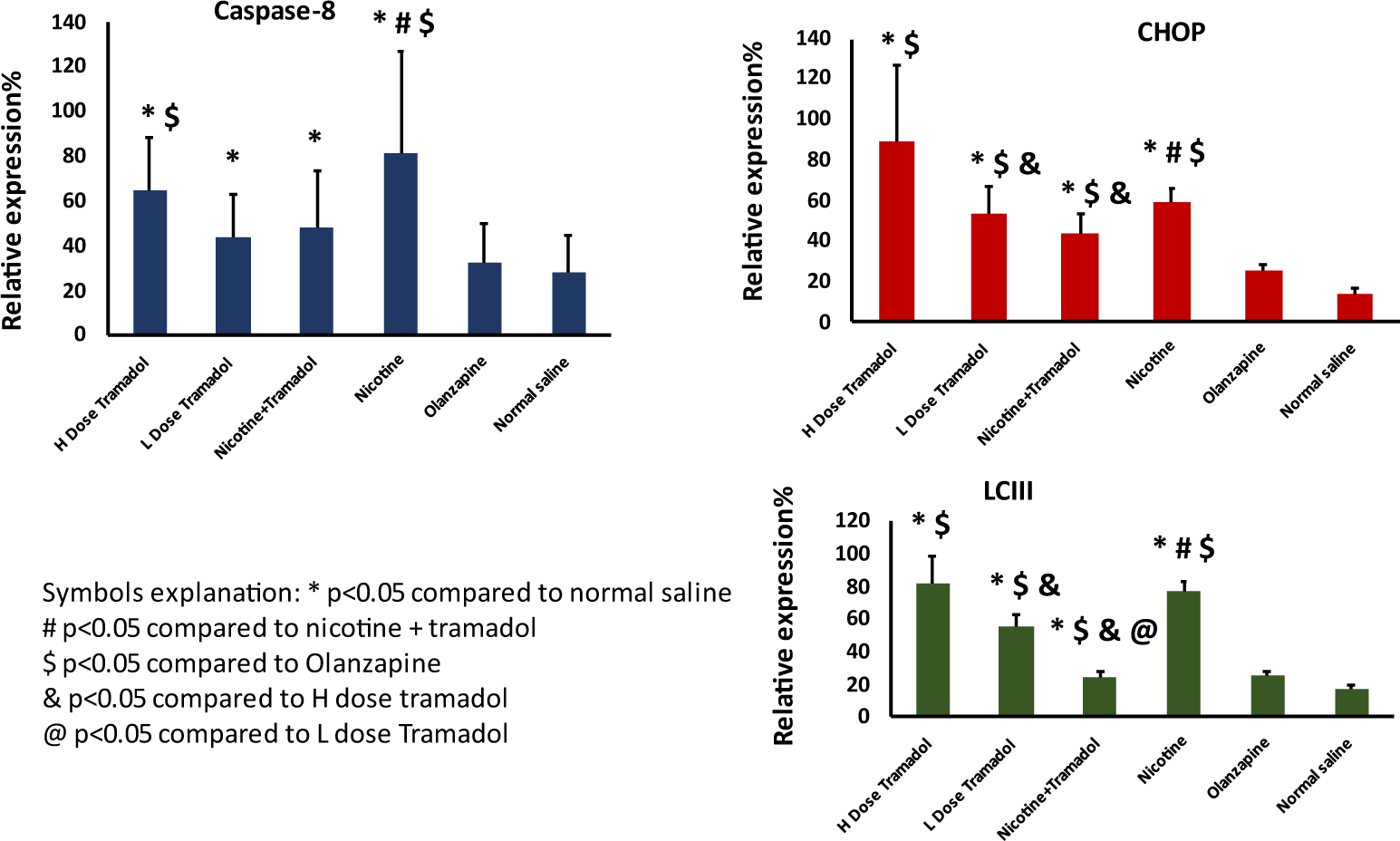
showed the relative expression of caspase 8, CHOP, and LCIII. The relative expressions of the 3 markers were demonstrated in the nicotine group. Relative lower expression was demonstrated in the nicotine/ tramadol combination

Figure 13 demonstrated the real-time gene expression of the CHOP gene in the animal rats. The results showed high gene expression in the control and the olanzapine groups. Relatively high expression was recorded in the high tramadol groups, and low expressions were expressed in the low tramadol groups and the combined tramadol/nicotine group. Moderate expression was noticed in nicotine groups.

**Figure (13):**
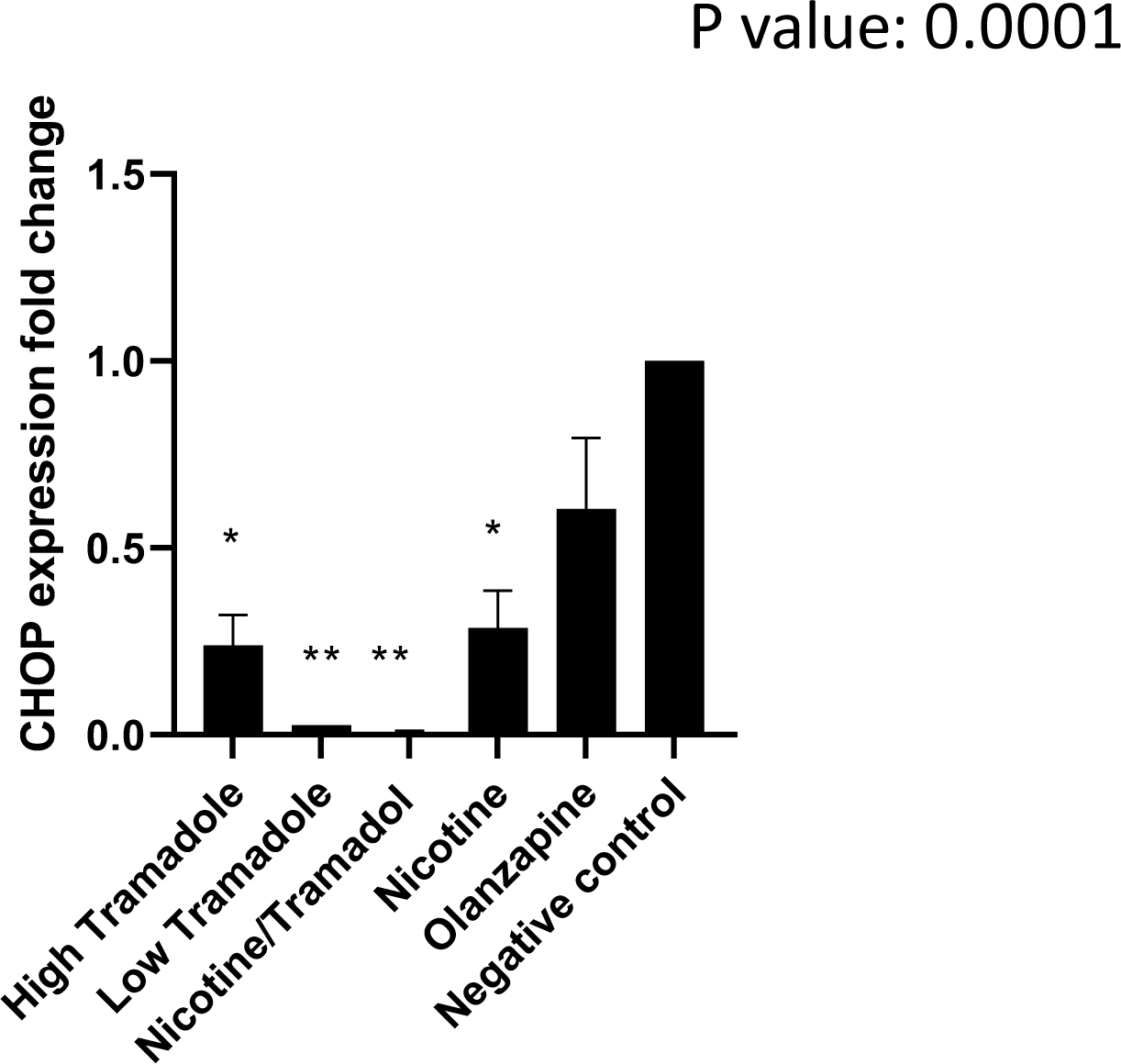
showed the relative gene expression of CHOP by real-time PCR. The highest expression was the negative control. The combined Nicotine/ tramadol expression was better than the nicotine group in rat brains.

Figure 14. showed the relative gene expression of BIP chaperone in rat livers. The figure revealed variable results which require interpretation. The highest expression was demonstrated in the low tramadol group with relatively high expression in the combined tramadol expression.

**Figure (14):**
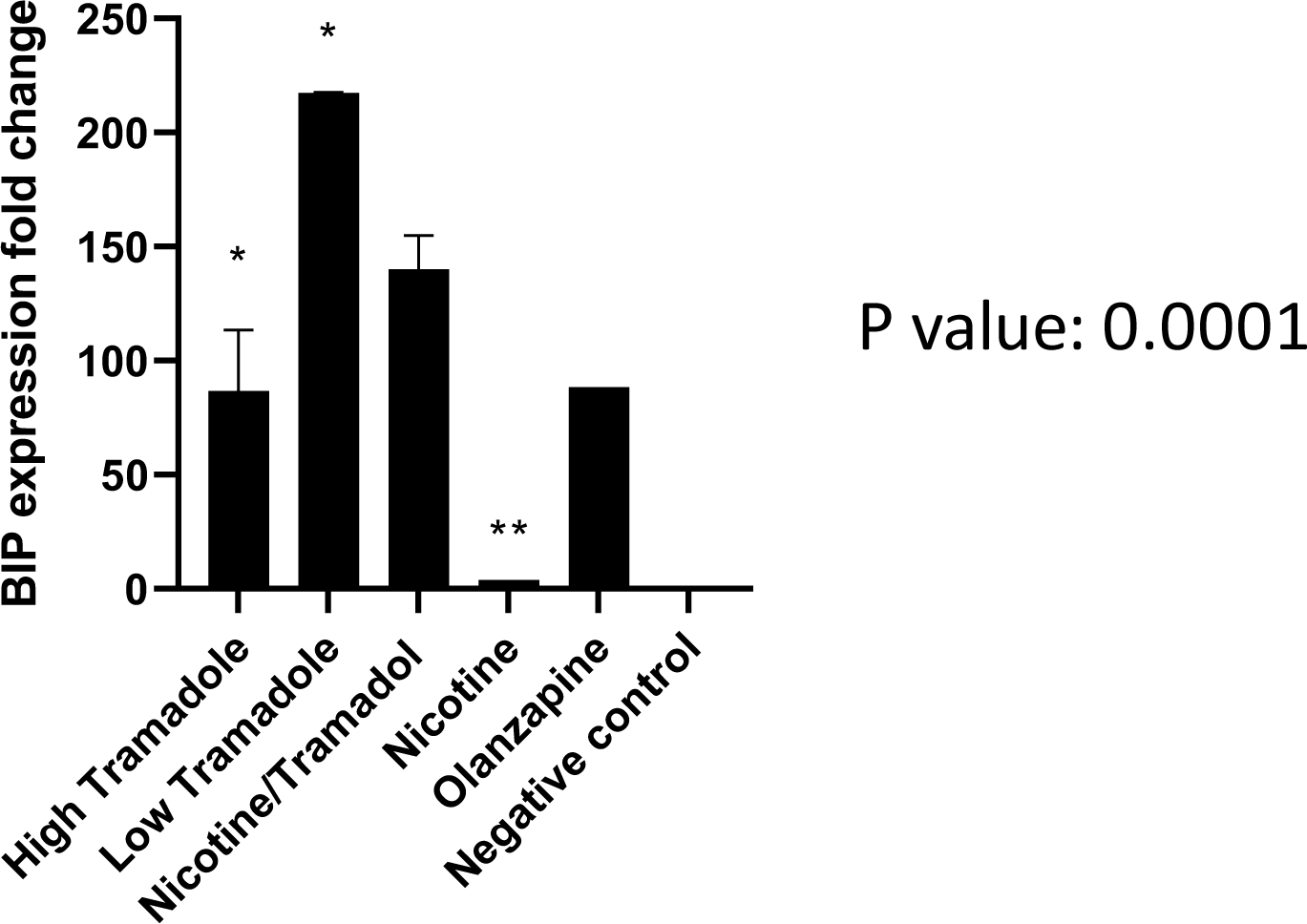
showed the relative expression of BIP protein in Rat liver. The gene expression, detected by real-time PCR, showed the highest expression in low tramadol and less expression in the combined nicotine/tramadol groups.

Figure 15 shows the relative serotonin blood level in the different 6 groups. The B component shows the relative blood level of androgen in different groups. As for serotonin, all groups demonstrated high serotonin blood levels in contrast to the negative control. The highest levels were the nicotine and nicotine/ tramadol combination. Also, the other groups, whether tramadol or olanzapine expressed high blood serotonin levels. On the other hand, no significant fluctuations were detected in the different groups of the androgen blood levels.

**Figure (15):**
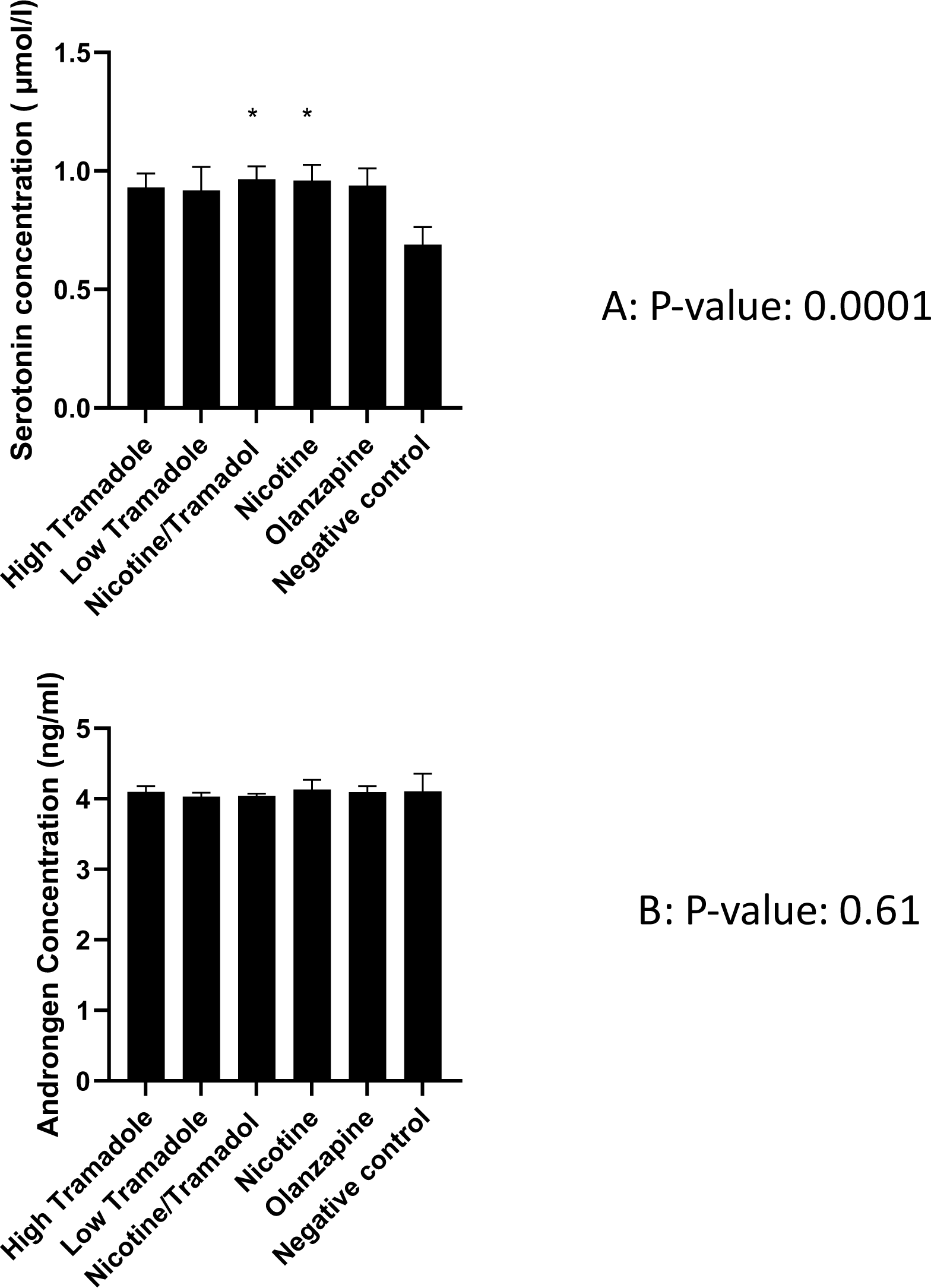
showed A: the serotonin, B: androgen levels in the animal’s blood. As for serotonin in the upper component of the figure, all groups showed statistically significant high expression of serotonin concerning the control. In the lower component of the figure. There was no statistically significant change in the androgen level in the rat blood in comparison with control.

Figure 16 illustrates the relative blood levels of dopamine blood levels of the animals in comparison to the saline groups. The nicotine group showed only decreased blood levels and serotonin. That reveals that nicotine increases serotonin and decreases dopamine in rat blood.

**Figure (16):**
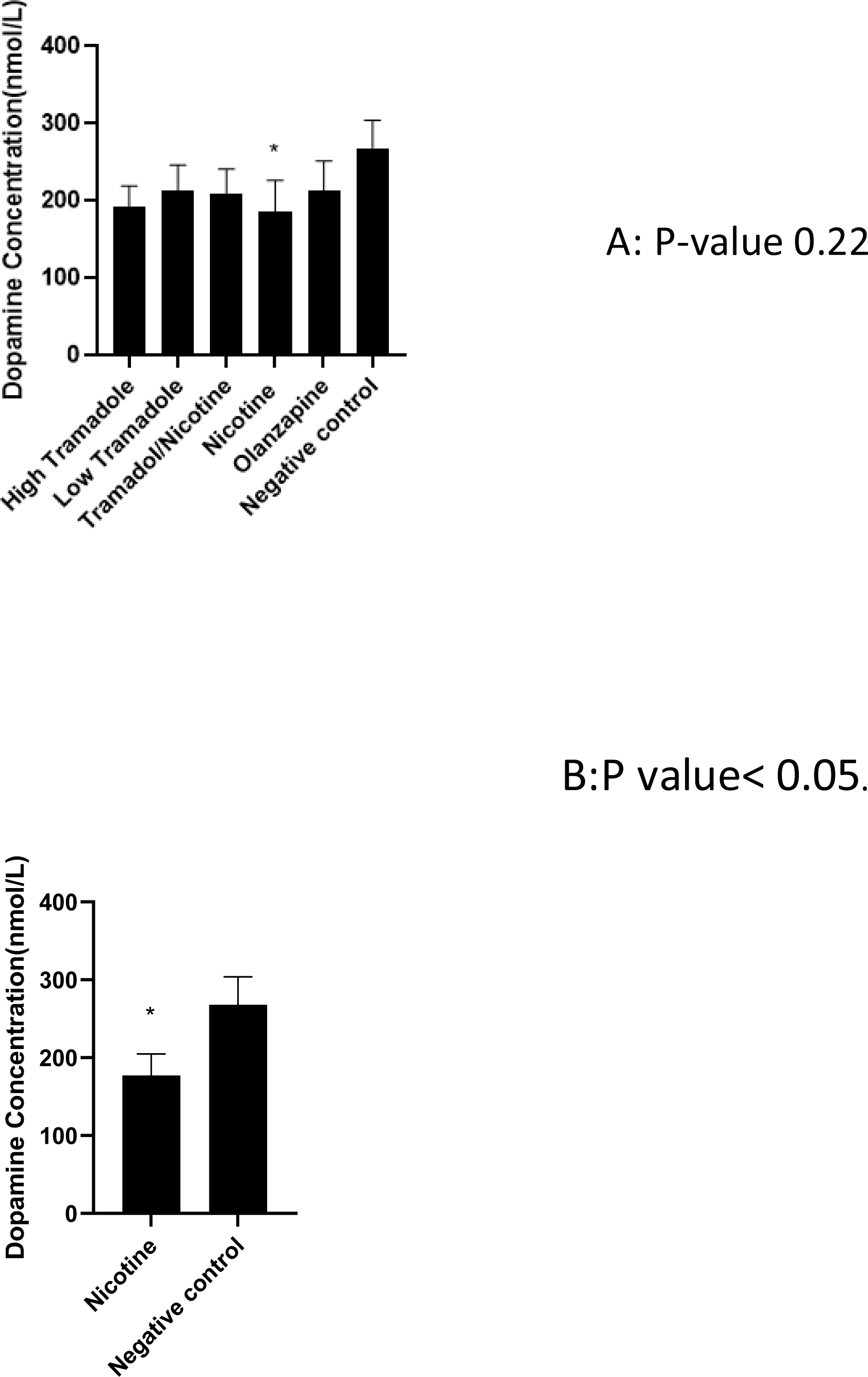
showed two components. A component expressed the animal blood levels of dopamine of the different groups in comparison with negative groups (high tramadol, low tramadol, combined tramadol nicotine, nicotine, and finally the negative control. There were no significant changes except in the relation between the nicotine group and negative control evaluated individually by t-test in the B component of the figure.

## Discussion

Prescription drug abuse, such as tramadol abuse, increases morbidity and death rates, while it also worsens the financial burden of indirect expenditures for health care, health promotion, surveillance, and dropped economic growth **(Silverman et al. 2022; Hassamal et al. 2018).** Prescription and controlled drugs are being used and abused more and more around the world, though the type of drug being abused may vary from country to country **(Sørensen et al. 2022; Goodarzi et al. 2011; Giraudon et al. 2013).**

As a centrally-acting analgesic, tramadol works through both opioid and nonopioid pathways **(Miotto et al. 2017)**. Its strong affinity for -opioid receptors and suppression of both norepinephrine and serotonin reuptake is what gives it its analgesic effects (Kaye 2015).

Smoking is a major risk factor for opioid addiction **(Rajabi et al. 2019)**, and using opioids and tobacco concurrently may improve perceived good effects and satisfaction with drug use, lessen withdrawal symptoms for both substances and serve as a substitute when one drug is unavailable **(David et al. 2013; David et al. 2014)**. Also, smoking enhances opioid usage **(Cooper et al. 2018)**. Despite the important overlap between nicotine and Tramadol, very few experimental research have looked into their combined impact. In this study, experimental animal work tracks the low toxic combination of tramadol/ nicotine on rats for 5 days per week for 3 weeks to explore the expected effects on the histology of (brain, liver, and testis), immunohistochemical staining, and western blotting for ER stress markers CHOP and BIB chaperones. Western blotting has been applied for Caspase 8 reflecting apoptosis and LC3 to detect autophagy **(Wang et al. 2016).** The work chose a lower toxic dose of tramadol for a shorter period focusing on ER stress. The experiment showed mild ER stress in the brain and demonstrated that both combined Nicotine/tramadol did not aggravate the condition.

Histological examination of the cortex demonstrated inflammatory response in the higher tramadol group and the combined tramadol /nicotine category although the tramadol-nicotine combination was healthy. On the other hand, the showing of the high tramadol group as the safest profile in hippocampus samples could be explained by the vasodilatation of the vessels in contrast to the other groups. All toxic groups showed injury to the animal’s white matter, including toxic olanzapine doses. The vasodilatation was reported to increase with the higher tramadol group. The CHOP immunohistochemistry showed positive staining in all groups. This finding can be augmented by the environmentalfactors where the place of the university is relatively 600 meters above the sea, and the climate is dry. Another explanation is the idea that ER stress is part of the physiological process in the neurons.

The p53 immunohistochemistry profile was the best in the nicotine animals and is in contrast to the combination animals. Regarding the low doses of smoking and mild tramadol abuse, the abusers look normal. As the study was applied for 3 weeks, little was known about chronic abusers.

The hippocampus immunohistochemical staining of CHOP showed improved Endoplasmic reticulum function if nicotine was added to tramadol. Minimal response to p53 immune stain was recorded in the hippocampus. It can be said that moderate tramadol use in conjunction with smoking is relatively safe for the hippocampus (**Xu et al. 2002)**.

As regards liver toxicity, all groups are relatively safe except mild hepatitis and vascular dilation in tramadol groups. These vascular phenomena may explain why abusers tend to use tramadol to improve sexual function according to an expert’s opinion **(Abdel-Hamid et al. 2016).**

Similar work demonstrated tramadol-induced ER stress in the adrenals which were associated with oxidative stress (Shalaby, Alabiad, and El Shaer 2020). Furthermore, more studies show that nicotine is a factor inducing ER stress (Wong, Holloway, and Hardy 2016a). In this work, the results reveal that both ER-inducing chemicals produce a better ER profile in combination. It suggests that nicotine attenuates the toxic response of tramadol at the level of the endoplasmic reticulum or at least did not worsen the condition. The positive effects of smoking on tramadol abuse have been seen in other experimental work that supports the current results (Azmy, Abd El Fattah, et al. 2018). The real-time PCR results of CHOP expression were similar but not typical in comparison with western blot results. Both western blot and qPCR showed the combined tramadol/nicotine group demonstrated less ER stress. The discrepancy between the protein expression detection by western blot and qPCR can be explained by the stability of the reference protein actin in comparison with GAPDH. The BIP protein and Caspase 8 expressions detected by western blot were a clear mirror of CHOP expression. Supporting the idea that moderate smoking in conjunction with moderate tramadol abuse appears stable. It is strongly recommended not to deal with both smoking and tramadol abuse in a separate management strategy. The Liver results showed more ER stress of both nicotine and tramadol than the control and confirmed that the combination was less problematic. It was demonstrated that the qPCR results varied from the final protein translation. It is suggested that the mRNA may be degraded early. Further research is needed to evaluate the validity of qPCR and the stability of mRNA in nicotine/tramadol models. The results of both high tramadol and low tramadol were variable regarding brain and Liver ER stress markers. It is suggested that the low margin change in the dose, the vasodilatation criterion, and the inter-animal response may explain the data.

This work shows that mild ER stress is suggested to promote the physiological functions of neurons and liver cells. The toxicity of tramadol, nicotine, or a combination was relatively mild. However, nicotine and tramadol at such low toxic doses for a relatively short period induced toxicity to the white matter. Higher tramadol doses showed vasodilation in both the brain and liver tissues Despite the promising histological benefits of mild ER stress, it is dangerous to depend on both smoking and tramadol abuse for longer periods. The ELIZA results expressed higher systemic levels of serotonin in animal blood. This hyper serotonin status will end in withdrawal and possible harmful adaptive neural response.

Olanzapine has been used as a positive control of ER stress. The present experiment showed that low-toxic doses of olanzapine promote mild ER stress. Furthermore, the histological profile was safe, however, the dose used in the course was associated with high blood serotonin despite that the drug is a well-known serotonin blocker. It seems that olanzapine exerts its mood stabilizer effect by a master key common hyper serotonin mechanism. On the other hand, this phenomenon may explain the withdrawal symptoms of the drug. As regards Tramadol, it is expected that it demonstrates high blood serotonin as it is a serotonin reuptake inhibitor. It has been recorded that tramadol abuser suffers from depression post-drug detoxification **(El-Hadidy and Helaly 2015)**. Other reports showed that post-tramadol abuse may end in psychosis (Rajabizadeh, Kheradmand, and Nasirian 2009; Sidana, Domun, and Arora 2019). In this study, it is recorded that excess serotonin may play a subsequent role in the development of both depression and psychosis. It is hypothesized that olanzapine users do not develop the same fate as tramadol ones because olanzapine blocks the serotonin receptor protecting it from the systemic hyper serotonin status. However, it is postulated to have atypical withdrawal symptoms. The present data show that the antidepressant effect can be induced by both blocking or stimulating the serotonin receptors. More research is promising to discover new drugs blocking serotonin receptors and increasing serotonin at the same time. It was hypothesized that serotonin may have a direct effect other than serotonin receptor signaling. It is important to notice that experimental work correlated high blood serotonin and the development of a metabolic syndrome or Diabetes with olanzapine toxicity. **(Erjavec et al. 2016)**. On evaluating the blood level of dopamine, the current study elucidates no systemic changes except in the nicotine group. Nicotine is known to increase dopamine release in the nucleus accumbens and substantia nigra. Interestingly, these findings varied in the inter-species responses **(Sziraki et al. 2001).** It was proposed that tramadol and nicotine would modify systemic dopamine as serotonin. However, nicotine lowered systemic dopamine without effects in other groups. Further work is needed to translate the results to humans

As regards testicular function, the different animal groups showed no significant changes in comparison with the negative control group. The ELIZA assay of androgen demonstrated a safe profile. Again, the reproductive results give a safe message about a moderate dose of tramadol in conjunction with smoking. This result conflicts with the results of many other robust studies. Tramadol, one of the most commonly abused medications affects spermatogenesis and messes with reproductive hormones **(Farag et al. 2018).** Abuse of tramadol is linked to low testosterone, hyperprolactinemia, and hypogonadotropic hypogonadism **(Attia et al. 2021).** However, the current model used more moderate abuse doses of tramadol than other studies for a relatively short period (Ghoneim et al. 2014). it is suggested that longer period high dose tramadol abusers may suffer infertility profile, unlike the current model.

The limitation of this study is the relatively short period of study. Also, it was needed to add a higher doses group to demonstrate evident apoptotic results. Further studies are needed to track higher doses of tramadol and nicotine for a longer period. More markers of ER stress can be tracked like PREK. Oxidative–inflammation response can be added to future work to study tramadol or nicotine in single or multiple experiments.

## Conclusion

The current study was able to explore the apparently healthy group abusing moderate dose tramadol and smoking which are misled by the potential favorable clinical outcome as long as they smoke and take tramadol regularly despite the potential threat of hyper serotonin impact on patients suffering from withdrawal. The present model showed that Tramadol, Nicotine, or the combination of two substances promotes mild ER stress. The combination is relatively safe with the potential of massive withdrawal syndrome like chronic depression. Further studies are needed to comprehend the role of serotonin and olanzapine in managing mood disorders.

## Acknowledgment

We are delighted to thank Lina Tahat for her supporting the statistics of this work. Many thanks to Amna El Rabie and Heba Al tammi for their technical support. We feel happy to thank Mr. Muhammad Yousef Abu Elrub for proofreading this work.

